# Tardigrade Dsup: Interactions with DNA and protection of cells from oxidative stress

**DOI:** 10.1101/2024.11.06.622393

**Authors:** Gavin Shuyang Ni, Hetian Su, Yingqi Zhu, Anshika Dhiman, Huan-Xiang Zhou, Wei Lin, Nan Hao

## Abstract

The remarkable capability of Tardigrade to survive under extreme conditions has been partially attributed to Dsup, an intrinsically disordered, highly positively charged protein. Dsup has been shown to bind to DNA in vitro, a property that has been associated with the capability of Dsup to exhibit stress-protective effects when expressed in mammalian cells. However, DNA binding of Dsup has not been visualized in living cells and expression of Dsup in different cell types was associated with either protective or detrimental effects. In addition, the effect of Dsup expression has not been clearly demonstrated at the organism level. Here we combined molecular dynamics (MD) simulations and fluorescence lifetime imaging microscopy (FLIM)-Förster resonance energy transfer (FRET) to interrogate Dsup-DNA interactions and demonstrated Dsup binding to DNA in living mammalian cells. Furthermore, Dsup expression in both HEK293T cells and yeast enhanced cell survival in the presence of hydrogen peroxide, suggesting that the presence of Dsup allows both mammalian and yeast cells to better cope with oxidative stress conditions. This study provides a better understanding of the property and functional role of Dsup and lays a foundation to explore new approaches to enhance stress resistance.

## Introduction

Tardigrades are a phylum of microorganisms that exhibit an incredible resistance to stress. They possess the ability to survive under extreme conditions such as intense heat and pressure and have been described as “practically unkillable”. They can survive in a wide variety of environments from scorching deserts to outer space, placing them in a category of animals known as extremophiles, a group of organisms known for their extreme survivability. This incredible resistive capability can be attributed to a combination of the damage suppressor protein and cryptobiosis.^1^ The damage suppressor protein, Dsup, has been found in the tardigrade *Ramazzottius varieornatus* and is partially responsible for the tardigrade’s resistive qualities.

Dsup is a 445-amino acid protein that is characterized by being largely unstructured and highly charged. It has a large number of charged residues, predominantly positively charged ones, with 59 lysine’s and 12 arginine’s, together with 29 aspartates and 19 glutamates accounting for 26.7% of the total amino acid composition. A computational study suggests that the intrinsically disordered nature of the protein enables Dsup to adopt different conformations to fit the shape of DNA.^2^ The electrostatic interactions facilitated by the charged residues and high protein flexibility lead to the formation of “a molecular aggregate” in which Dsup shields DNA. Dsup was found to be localized in the nucleus and shown to prefer to bind to nucleosomes as opposed to free DNA *in vitro*.^3^ Dsup has also been shown to be localized to the nucleus and bind to chromatin in yeast.^4^ However, Dsup binding to DNA has not been directly visualized in living systems. Functionally, Dsup has been demonstrated to protect the DNA from damage. For example, Dsup, when expressed in cultured HEK293T cells, suppressed X-ray induced DNA damage by 40%.^5^ However, in a different study, Dsup expression in cortical neurons resulted in the formation of DNA double-stranded breaks, suggesting that Dsup may promote DNA damage in this system.^6^ Thus, the binding preference and the functional effects of Dsup have not been well understood.

Oxygen, a crucial component for the survival of many organisms, has the tendency to produce reactive oxygen species (ROS). Common types of reactive oxygen species are the hydroxyl radical, the superoxide radical, and hydrogen peroxide (H_2_O_2_). These types of ROS are typically due to errors in the electron transport chain during cellular respiration,^7^ and can also be induced by environmental stimuli such as exposure to ultraviolet radiation and toxic heavy metals—cadmium or lead.^8^ ROS causes oxidative stress to the cells, catalyzes the aging process, and is linked to the onset and progression of various diseases such as cancer, Parkinson’s, Alzheimer’s, and cardiovascular diseases.^9^ ROS can cause many different forms of damage to DNA, including strand breaks, base substitution, and protein-DNA cross-links. These damages result in either programmed cell death or compromised DNA functionality, which could lead to the development of cancer.^7^ The effects of Dsup on protecting cultured cells and whole organisms other than Tardigrades from ROS-mediated DNA damages await further investigation.

In this study, we first applied MD simulations and FLIM-FRET to probe Dsup-DNA interactions in silico and in living mammalian cells, respectively. Furthermore, using HEK293T cells as well as a yeast strain expressing Dsup under the control of doxycycline, we tested the effect of Dusp on these cells after exposure to H_2_O_2_. Our study sheds light on the damage suppressor protein’s ability to protect DNA in different living systems, bringing us one step closer to a possible solution for oxidative stress linked diseases such as cancer and oxidative stress induced aging.

## Results

### MD simulations of Dsup-DNA interactions

The DNA-binding region of Dsup was isolated to the C-terminal ∼90 residues,^3^ which are rich in basic residues. To characterize the Dsup-DNA interactions in atomic details, we performed MD simulations of three copies of the C-terminal region (residues 361-445; 6 R, 14 K, no D or E) in complex with a 35-base pair DNA. Similar to an arginine-rich intrinsically disordered protein protamine,^10^ the Dsup chains quickly collapsed around the DNA; Arg and Lys residues formed both salt bridges with both the DNA backbone phosphates and hydrogen bonds with the nucleobases (Fig. 1a). Arg in particular frequently hydrogen bonds with two or more bases (Fig. 1a, zoomed view). On average, 35.1 residues of the Dsup chains interacted with phosphates and 10.6 residues interacted with bases; each Dusp-interacting phosphate engaged with 1.2 Dsup residues while each DNA-interacting residue engaged with 1.7 bases (Fig. 1b).

**Figure 1.**
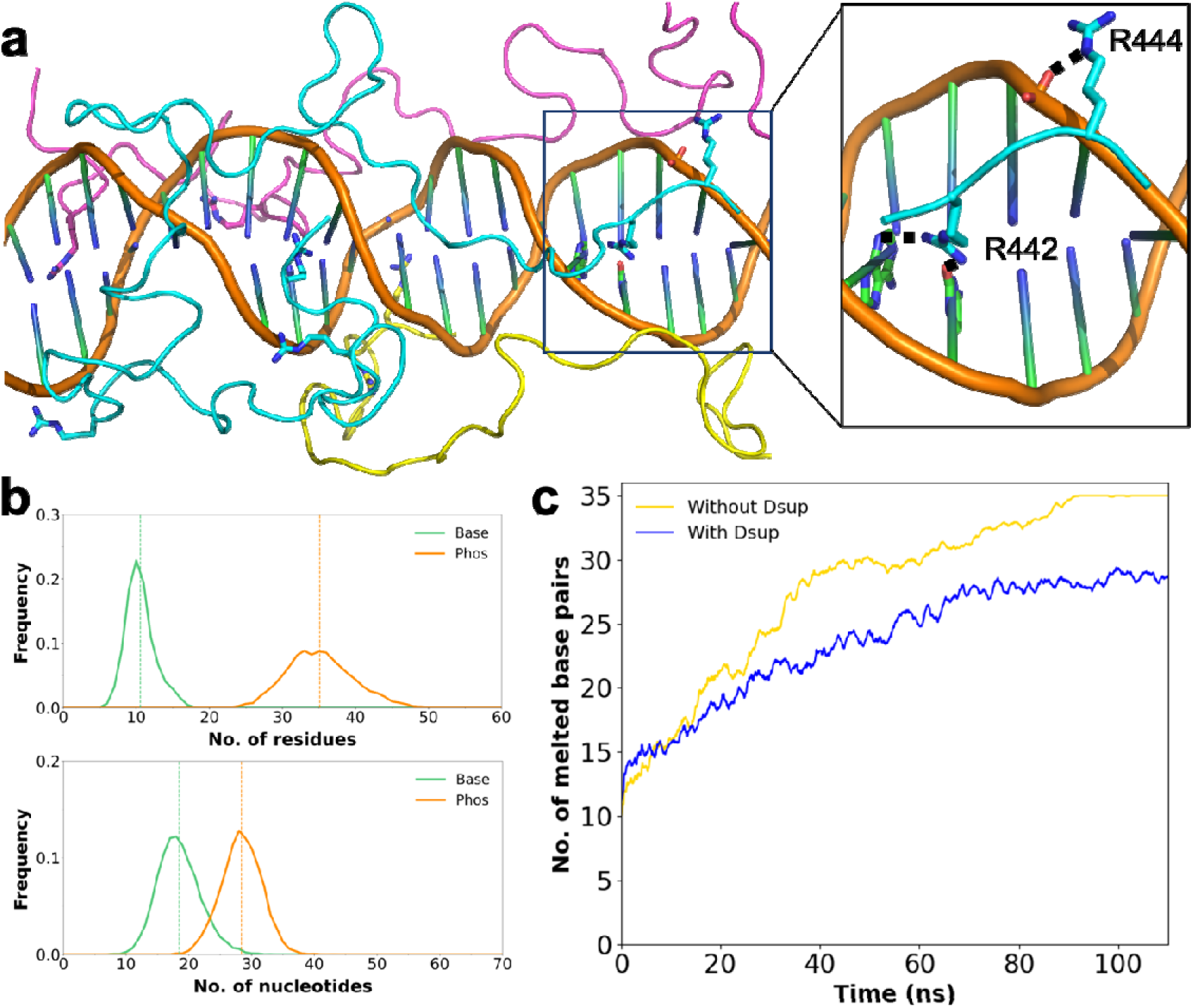
MD simulations of Dsup’s interactions with DNA and protective effect on DNA melting. **a.** A snapshot from the simulations at 300 K. Inset: hydrogen bonding of R442 with two nucleobases and salt bridge of R444 with a backbone phosphate. **b.** Distribution of the number of Dsup residues interacting with DNA bases or phosphates. **c.** Distribution of the number of bases or phosphates interacting with Dsup residues. d. The number of melted base pairs in the absence and presence of Dsup in simulations at 500 K. Time traces were smoothed as the running average in a 11-ns window.

To directly demonstrate the protective effect of Dsup, we melted the DNA in simulations at 500 K in the absence and presence of Dsup. Without Dsup, the DNA melted completely in 92 ns (Fig. 1c). In contrast, with the three Dsup chains initially bound, the DNA melted much more slowly; 8 base pairs remained intact after 92 ns of simulations.

### FLIM-FRET demonstrates Dsup binding to DNA in living mammalian cells

To directly probe Dsup-DNA interactions in living mammalian cells, we set out to develop an assay based on FRET, a well-known spectroscopic ruler for monitoring molecular interactions in live cells. In combination with FLIM, FRET can be quantitatively read out through measuring the fluorescence lifetime changes of donor fluorophore. FLIM-FRET has been used as a reliable tool to detect the *in situ* interaction between protein-DNA in the cell nucleus.^11^ To establish a FLIM-FRET assay for Dsup-DNA interaction, we selected enhanced green fluorescent protein (EGFP) as the FRET donor to fuse to the C-terminal of Dsup protein and SYTO™ 59 Red Fluorescent Nucleic Acid Stain as the FRET acceptor to stain DNA. When Dsup binds to DNA, EGFP along with Dsup was expected to be in close proximity to SYTO™ 59 on DNA, leading to a shortened fluorescence lifetime of EGFP.

To first establish the DNA-protein assay, we performed a positive control experiment using histone 2B (H2B)-EGFP which is known to be tightly wrapped by chromatin DNA. HEK293T cells transiently expressing H2B-EGFP were stained with SYTO™ 59, followed by FLIM-FRET imaging. Upon binding to DNA, the red fluorescence from SYTO™ 59 was mainly detected in the nuclear area. H2B-EGFP was co-localized in the nuclei with SYTO™ 59 stained DNA (Fig. 2a), and significantly short lifetime from H2B-EGFP was observed (τ = 1.78 ± 0.02 ns, n = 26 cells) compared to H2B-EGFP only cells (τ = 2.48 ± 0.02 ns, n = 17 cells) (Fig. 2b). Next, we stained HEK293T cells transiently expressing Dsup-EGFP with SYTO™ 59. As shown in Figure 2a, we observed green fluorescence in the nuclei indicating the subcellular location of Dsup. We then performed FLIM-FRET measurements and observed significantly shorter donor fluorescence lifetime within nuclei from Dsup-EGFP cells stained with SYTO™ 59 (τ = 1.79 ± 0.02 ns, n = 24 cells) compared with Dsup-EGFP cells without SYTO™ 59 staining (τ = 2.51 ± 0.01 ns, n = 19 cells) (Fig. 2c). To further rule out the possibility of non-specific intermolecular FRET, we performed a negative control experiment with nuclear localization signal (NLS)-tagged EGFP expressed in the nucleus. From images, EGFP-NLS was largely directed to the nuclear plasm co-localizing with SYTO™ 59 stained DNA (Fig. 2a). As expected, EGFP-NLS expressing cells showed no difference in the presence of SYTO™ 59 staining (τ = 2.45 ± 0.01 ns, n = 29 cells) compared to EGFP-NLS only cells (τ = 2.45 ± 0.01 ns, n = 24 cells) (Fig. 2d). Together, these results provide direct evidence for association between Dsup and DNA within the nuclei of living mammalian cells.

**Figure 2.**
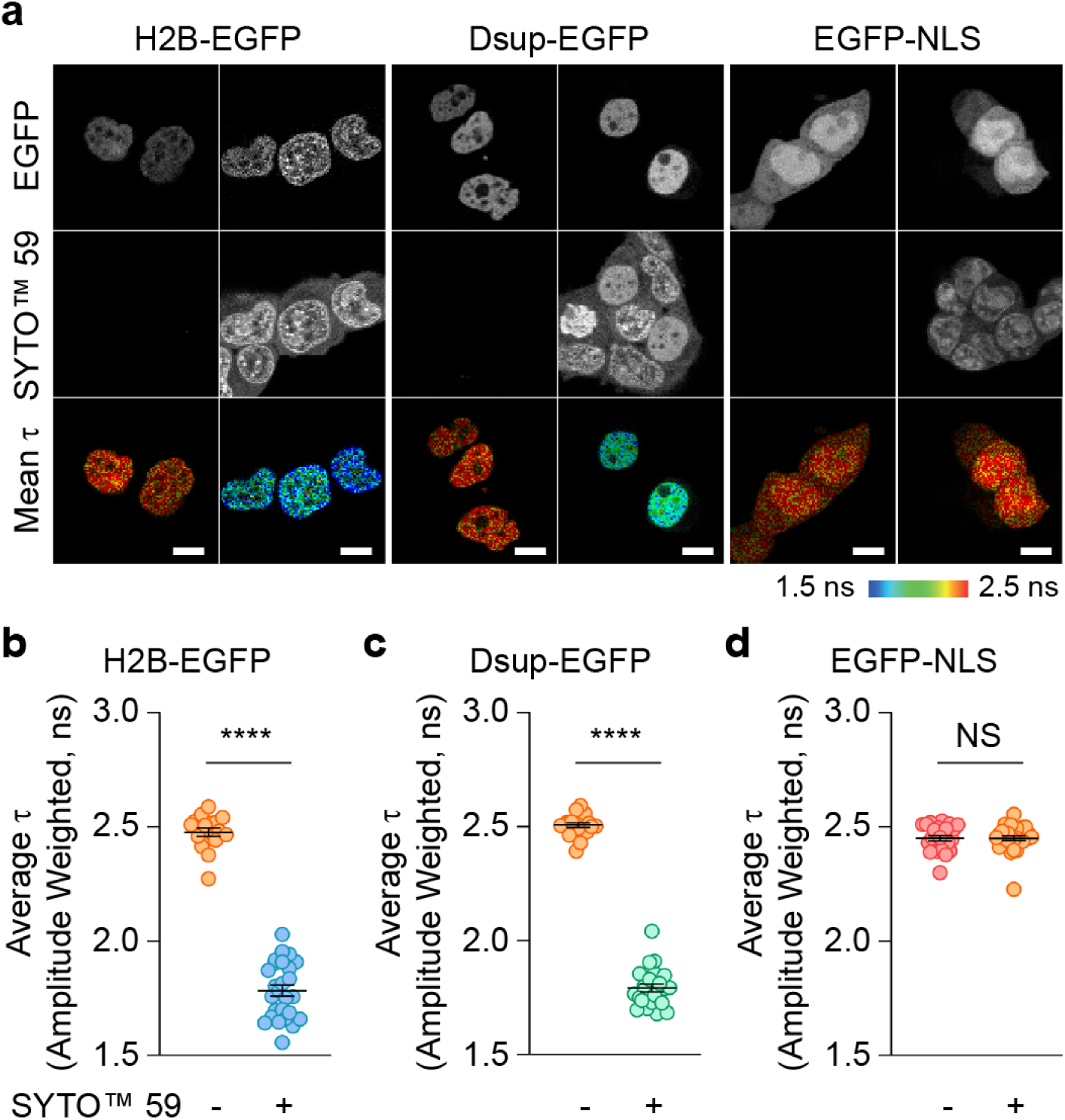
FLIM-FRET demonstrates Dsup binding to DNA in HEK293T cells. **a.** Representative FLIM-FRET images of H2B-EGFP (left), Dsup-EGFP (middle) and EGFP-NLS (right). Cells were stained with or without SYTO^TM^ 59. From the top to bottom are shown the EGFP fluorescence intensity images, SYTO^TM^ 59 fluorescence intensity images, and fast FLIM maps. The color-coded scale corresponds to the fluorescence lifetime. Scale bar: 10 µm. **b-d.** Quantification of the average fluorescence lifetime (τ) of single nuclei of H2B-EGFP (**b**), Dsup-EGFP (**c**) and EGFP-NLS (**d**) expressing cells with or without SYTO^TM^ 59 staining. Data from 3 independent experiments. Statistical analyses were performed using unpaired two-tailed Student’s *t*-test. \*\*\*\**P* < 0.0001. NS, not significant. Data are mean ± s.e.m.

### Dsup expression enhances cell survival in the presence of hydrogen peroxide

High levels of hydrogen peroxide are known to induce oxidative stress and cause cell death through a variety of mechanisms including hydrogen peroxide-induced DNA damage.^12, 13^ We set out to investigate the effect of Dsup expression on HEK293T cell survival exposed to H_2_O_2_. Overexpression of Dsup over extended time has been reported to increase cell death,^6^ we thus generated a HEK293T stable cell line expressing Dsup-EGFP, along with a negative control HEK293T line expressing only EGFP, both regulated by a Tet-on promoter and doxycycline.^14^ As demonstrated in Figure 3a, live-cell imaging revealed the nuclear localization of Dsup and the nucleocytoplasmic localization of the EGFP control upon doxycycline (12.5 ng mL^−1^) induction, verifying the correct subcellular localization of these proteins.

**Figure 3.**
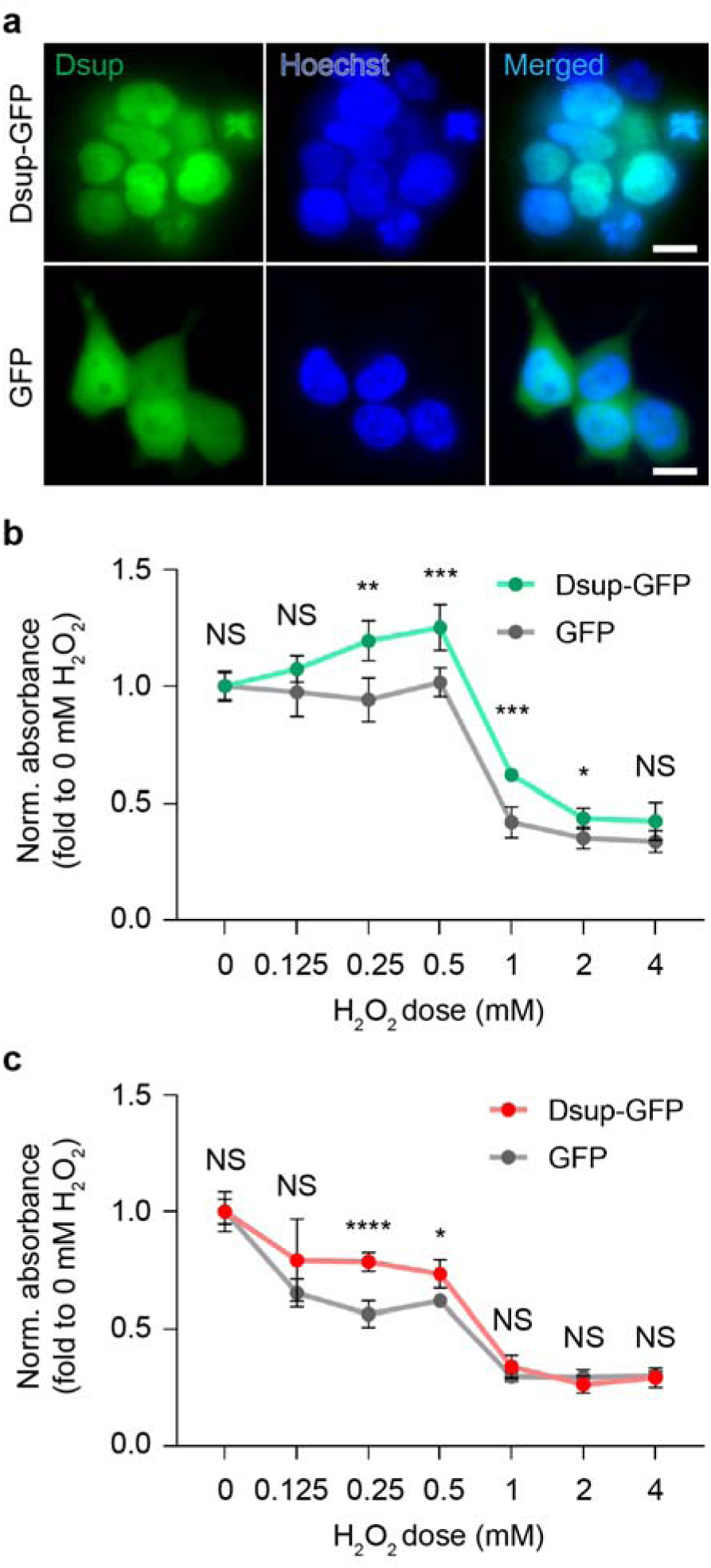
Dsup expression enhances cell survival in the presence of hydrogen peroxide. **a.** Representative images of induced expression of Dsup-EGFP (upper) and EGFP (lower) by doxycycline in HEK293T cells. The nuclei were stained with Hoechst 33342. Scale bar: 10 µm. **b-c.** Normalized absorbance of formazan dye converted by dehydrogenases from viable cells expressing Dsup-EGFP or EGFP. H_2_O_2_ doses from 0 to 4 mM were applied for 24 hours (**b**) and 48 hours (**c**). Data from 3 independent experiments. Statistical analyses were performed using unpaired two-tailed Student’s *t*-tests. \*\**P* = 0.00126 (24 hr, 0.25 mM), \*\*\**P* = 0.00092 (24 hr, 0.5 mM), \*\*\**P* = 0.00013 (24 hr, 1 mM), \**P* = 0.0152 (24 hr, 2 mM), \**P* = 0.0219 (48 hr, 0.5 mM), \*\*\*\**P* < 0.0001. NS, not significant. Data are mean ± s.d.

To explore Dsup’s role in protecting genomic DNA against H_2_O_2_, we employed the Cell Counting Kit-8 (CCK-8) assay, a widely used colorimetric technique for assessing cell proliferation. Cells were exposed to varying concentrations of H_2_O_2_ for 24 hours and dehydrogenase activities in cells were measured and compared. Notably, Dsup expression increased cell viability compared to the EGFP control under conditions from 0.25 mM to 2 mM H_2_O_2_ (Figure 3b). After a 48-hour treatment with H_2_O_2_, Dsup expression increased cell viability compared to the EGFP control at 0.25 mM and 0.5 mM HLOL, while no significant differences were observed at either lower or higher H_2_O_2_ concentrations (Figure 3c). The null effect at lower H_2_O_2_ can be explained by a lack of H_2_O_2_ damage, whereas that at higher H_2_O_2_ can be attributed to an overwhelming level of H_2_O_2_ damage. These findings suggest that Dsup expression in mammalian cells improves cell survival under oxidative stress, particularly in the presence of reactive oxygen species like hydrogen peroxide.

### Effects of the Dsup protein from Tardigrade on stress-resistance of yeast

To further evaluate the effects of Dsup on another organism, we turned to budding yeast *Saccharomyces cerevisiae* and generated a genetically engineered budding yeast strain where Dsup expression is under doxycycline control. Doxycycline was used to control the Tet-on promoter and express the Dsup tagged on the N-terminus by a red fluorescent protein mRuby^15^ to track the expression level of Dsup. As shown in Figure 4a, we observed red fluorescence in the nuclei, which colocalized with the Nhp4a-iRFP nuclear marker. We analyzed the expression of Dsup by tracking the red fluorescence intensity as a function of time in yeast cells with Dsup expression induced by 500 nM doxycycline, compared with no doxycycline control. The expression of Dsup in cells increased with time in the presence of doxycycline, whereas there was generally no Dsup expression in the absence of doxycycline, showing low leakage of the dox-inducible promoter system (Figure 4b).

**Figure 4.**
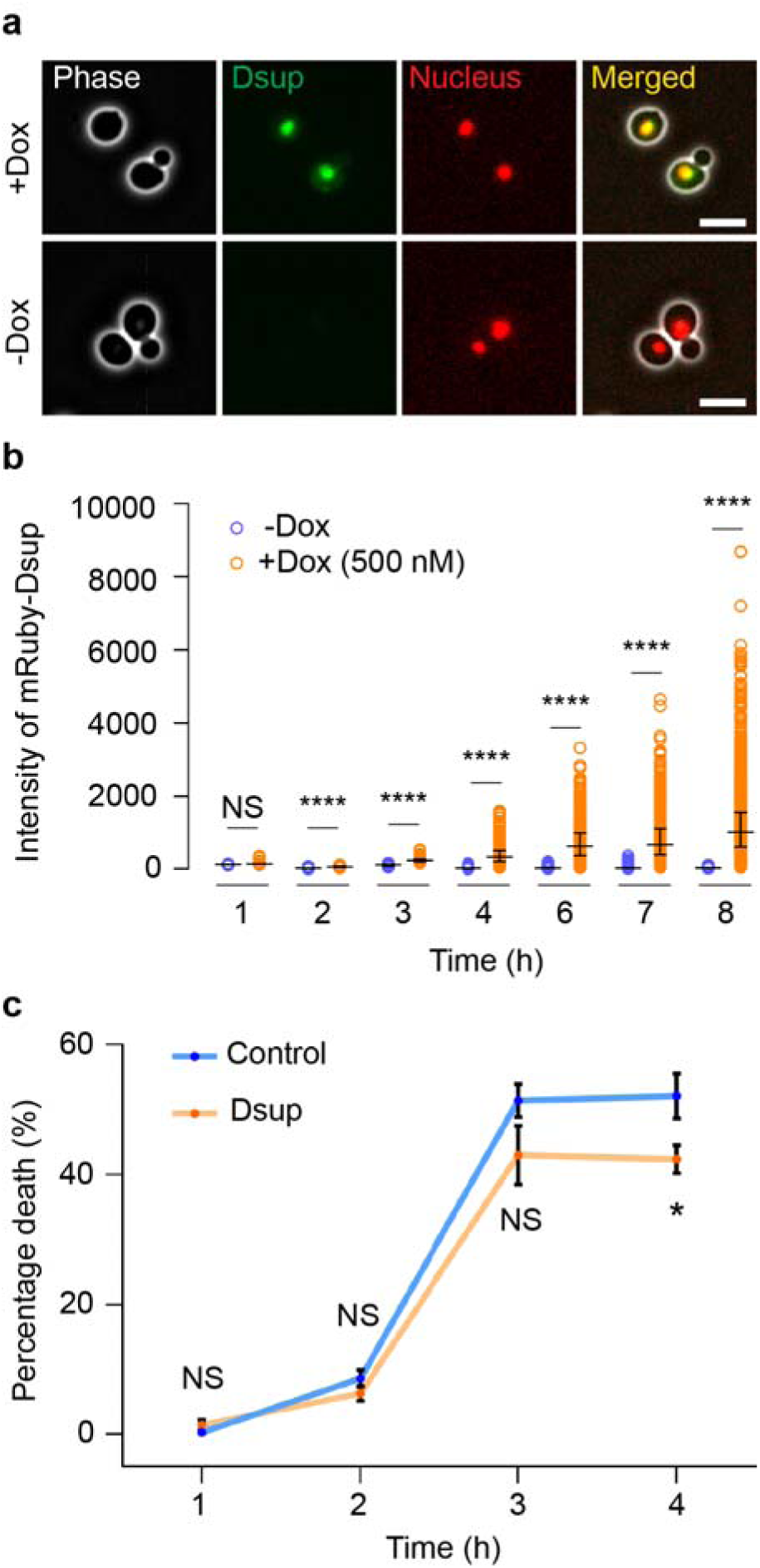
Effects of the Dsup protein on H_2_O_2_ stress-resistance of yeast. **a.** Representative images of induced expression of mRuby-Dsup in yeast. The nuclei were indicated by Nhp4a-iRFP nuclear marker. Scale bar: 5 µm **b.** Quantification of time-course expression of mRuby-Dsup induced by doxycycline (Dox) or not. Statistical analyses were performed using two-sided Student’s *t*-test (N = 5). \*\*\*\**P* < 0.0001. NS, not significant. Data are median with interquartile range. **c.** Cell survival assay by PI dye staining. Statistical analyses were performed using the Mann-Whitney U test (N = 9). \**P* = 0.0157. NS, not significant. Data are mean ± s.e.m.

We then treated yeast cells with H_2_O_2_ in liquid culture and used propidium iodide (PI) staining to identify severely damaged cells. In order to avoid effects of fluorescent protein tagging on Dsup’s function, we used a yeast strain containing untagged, dox-inducible Dsup. As shown in Figure 4c, three and four hours after H_2_O_2_ (2 µM) addition, cells with Dsup induction exhibited 43.0 ± 4.5% and 42.4 ± 2.2% of cell death, while 51.4 ± 2.6% and 52.1 ± 3.5% of cell death were exhibited by those without Dsup induction (Figure 4c). This is due to more cells being severely damaged and stained by PI in non-induction cells. These results indicate that after four hours of exposure to hydrogen peroxide, the Dsup-expressing cells started to exhibit statistically significant lower cell death, than cells without Dsup. Thus, the expression of Dsup in yeast is beneficial for its survival under oxidative stresses caused by hydrogen peroxide.

## Discussion

Buildup of excess ROS places oxidative stress on cells, altering base pairs and causing double-stranded breaks, leading to impaired DNA functionality among other detrimental effects. Oxidative stress and the resulting damage can catalyze the aging process as well as cause the development of multiple diseases. Identification of factors that confer resistance to oxidative stress induced damage could aid in the slowing of the aging process and the prevention of serious diseases. The investigation of stress-resistance is therefore important for study of aging and for addressing unmet needs of people affected by aging related disorders. Tardigrades, otherwise known as water bears, are near-microscopic animals well known for their incredible resistive capabilities under extreme conditions such as intense radiation and in outer space. The tardigrade Dsup protein was suggested to protect DNA from various stresses and damages. However, Dsup expression in cortical neurons was also reported to result in the formation of DNA double-stranded breaks, suggesting that Dsup may promote DNA damage in some systems.^6^ In this study, we investigated the interactions between Dsup and DNA both in silico and in living cells and probed the role of Dsup in protecting mammalian and yeast cells from H_2_O_2_-induced toxicity.

To provide a mechanistic understanding of the protective effects of Dsup, we performed MD simulations of DNA binding with the C-terminal disordered region of Dsup. The simulations showed that Dsup chains wrap around DNA, with Arg and Lys interacting with the DNA backbone phosphates and nucleobases via salt bridges and hydrogen bonds, respectively. Through these interactions, Dsup may provide a physical barrier to prevent ROS-induced double-stranded breaks and damage on DNA base pairs. Additionally, Dsup may keep DNA from structural changes that are susceptible to damage. Indeed, our simulations at 500 K revealed that Dsup slows down the melting of DNA. The resulting reduction in disrupted base pairs (Fig. 1c) appears qualitatively similar to the reduction in cell death (Fig. 4c). Although we caution against drawing a direct link between these two effects, it stands to reason that strand separation or other more subtle and transient structural changes can expose nucleobases and thereby make them more vulnerable to ROS-induced reactions.

Dsup was shown to bind to both free DNA and nucleosome *in vitro*, with enhanced binding towards nucleosome.^3^ Our data clearly demonstrates Dsup-DNA interactions in the nuclei of living mammalian cells, which complements to the *in vitro* study. Dsup-expressing mammalian cells can be readily stained by the red fluorescent nucleic acid dye, suggesting that the small molecule binding to DNA and Dsup binding to DNA are not interfering with each other. The FLIM-FRET-based imaging method for studying biomolecular interactions offers quantitative and sensitive readout. Although widely used as a method for detecting protein-protein interactions, there were fewer examples of live-cell FLIM-FRET studies on interactions between DNA and DNA-binding proteins. Our study demonstrated that the GFP-SYTO based FLIM-FRET assay is an efficient method for detecting Dsup-DNA interactions *in situ* and should be readily extensible to other systems.

We further showed Dsup expression protects HEK293T cells from oxidative stress, consistent with the findings reported by Hashimoto *et al*.^5^ However, the observed toxicity resulting from high-level over-expression of Dsup could underlie the detrimental effects observed by Diaz Escarcega *et al*.^6^ Budding yeast is a commonly used model organism due to its genetical tractability. Importantly, budding yeast retains many of the core eukaryotic cellular processes including elements of programmed cell death (PCD) such as apoptosis. Similar to mammalian cells, ROS plays a major role in PCD pathways in yeast.^7^ Also, aging and oxidative stress are easily tractable in yeast due to its short lifetime and rapid division. Prior to our study, the effects of Dsup in protecting yeast from oxidative stress have been reported.^4^ In agreement with this work, our study, by measuring the survival of yeast cells in liquid cultures, showed Dsup expression could protect yeast from hydrogen peroxide induced stress and damage at micromolar concentration level. Together, we demonstrated that the function of Dsup could be universal across different species. This study sheds light on the possibility of using yeast as a simple model to further investigate dynamic control of Dsup as a possible solution for oxidative stress linked, age-related diseases such as cancer, Parkinson’s disease and Alzheimer’s disease.

## Acknowledgments

The authors thank all members of the Hao lab for various technical help and discussion of the manuscript. This work is supported by NIH grants R01AG056440, R01GM144595, R01AG068112, R01AG086348, and R35GM118091. This work is also supported by the UCSD Microscopy Core (P30 NS047101).

## Author contributions

G.N. and N.H. conceived of the project. G.N., H.S., Y.Z. and W.L. performed the experiments and analyzed the data. A.D. performed MD simulations. H.X.Z., W.L., and N.H. supervised the research. G.N., H.S., Y.Z., A.D., H.X.Z. and W.L. wrote the manuscript. Every author read and agreed on the final manuscript.

## Methods

### Materials

Hydrogen peroxide solution (30%, H325-100) was ordered from Fisher Chemical, Doxycycline hyclate (Sigma-Aldrich, P5207), propidium iodide (Sigma-Aldrich, P4170) were reconstituted in DMSO (Sigma-Aldrich) as 1000× stock solutions. UTP (Alfa, 98%+, J63427) was reconstituted in ultrapure water (MilliQ). Lipofectamine 2000 (11668019) and SYTO™ 59 (S11341) were purchased from Thermo Fisher. Cell Counting Kit-8 (GK10001) was purchased from GLPBIO.

### Plasmids

H2B-EGFP-N1 was ordered from Addgene (#11680). For EGFP-NLS construct, EGFP-NLS was subcloned to pcDNA3 vector between *Bam*HI and *Xba*I restriction sites. For Dsup-EGFP construct, the DNA fragment encoding Dsup was amplified using primers 5’- TCA AGC TTC GAA TTC TGC AGT CGA CAT GGC ATC CAC ACA CCA ATC - 3’ and 5’ – TCA CCA TGG TGG CGA CCG GTG GAT CCT TCC TCT TCC GTC CTC CAG - 3’, and replaced H2B in H2B-EGFP-N1 by Gibson Assembly. pLenti-CMV-rtTA3-Blast (#26429), psPAX2 (#12260) and pMD2.G (#12259) were ordered from Addgene. pLenti-TetO-EGFP-Puro construct was generated previously.^16^ For pLenti-TetO-Dsup-EGFP-Puro construct, the DNA fragment encoding Dsup was amplified using primers 5’- ATT CAA GGC CTC TCG AGC CTC TAG AAT GGC ATC CAC ACA CC - 3’ and 5’ - GCT CAC CAT GGT GGC GAC CGG TGG ATC CTT CCT CTT CCG - 3’ were cloned into pLenti-TetO-EGFP-Puro construct fusing to 5’- terminus of EGFP. For Tet-on mRuby-Dsup and Tet-on Dsup constructs, A Moclo Leu2 cassette with homology arms to the upstream and downstream of yeast Leu2 was synthesized, a ScRPL18B promoter-rtTA3-ScADH1 terminator fragment was synthesized on pYTK95 backbone, and a pTetO7-Dsup-tENO2, or a pTetO7-mRuby-Dsup-tENO2, fragment was synthesized on pYTK95 backbone. These constructs were digested with BsmBI and assembled with Golden Gate Assembly to form the full doxycycline-inducible construct with (NHB1086) or without (NHB1054) mRuby tagging.

### Cell culture and transfection

HEK293T (ATCC, CRL-3216) cells were cultured in Dulbecco’s modified Eagle medium (DMEM) (Gibco) containing 4.5 g l^−1^ glucose, 10% (v/v) fetal bovine serum (FBS) (Sigma), and 1% (v/v) penicillin-streptomycin (Pen-Strep) (Sigma-Aldrich). All cells were maintained at 37 °C in a humidified atmosphere of 5% CO_2_. For imaging experiments, cells were plated onto sterile 35-mm glass-bottomed dishes and grown to 50–70% confluence. Transient transfection was performed using PolyJet following manufacturer’s protocols, after which cells were cultured for an additional 24 h.

### HEK293T stable cell line generation

HEK293T stable cell lines were generated using lentiviral plasmid vectors (plenti-TetO-Dsup-EGFP, plenti-TetO-EGFP, and plenti-rtTA) by dual-infection. To package the lentivirus containing TetO-Dsup-EGFP, TetO-EGFP, and rtTA, each pLenti-construct was mixed with psPAX2 and pMD2.G-VSV-g (3:2:1) and co-transfected into HEK293T cells. The medium containing packaged lentivirus was collected after 24 h and directly added to HEK293T in 6 cm cultured dish to generate rtTA + TetO-EGFP and rtTA + TetO-Dsup-EGFP dual-stable cells. After 24 h, cells were diluted and plated in 10 cm cell culture dishes and then treated with Puromycin (1LµgLmL^−1^) and Blasticidin (10 µgLmL^−1^) to select transduced cells. For cell maintenance, cells were cultured in the medium with Puromycin (0.5LµgLmL^−1^) and Blasticidin (5 µgLmL^−1^). Doxycycline will induce the protein expression only when both rtTA and TetO reporters integrated to cellular genome.

### CCK-8 colorimetric assay

Cells were seeded in a 96-well plate at a density of 15k cells per well in 100 μL of culture medium and incubated at 37°C with 5% CO_2_ for 24 hours. Various concentrations of H_2_O_2_ and 12.5ng mL^−1^ of Doxycycline were added to the cells for 48 hours. Following this, 10 μL of CCK-8 solution was added to each well, and the plate was incubated for an additional 1 hr. After gently mixing the plate on an orbital shaker, the absorbance was measured at 450 nm using a microplate reader.

### Yeast cell culture

Standard methods for the growth, maintenance and transformation of yeast and bacteria and for manipulation of DNA were used throughout. The yeast strains used in this study were generated from BY4741 (MATa his3D1 leu2D0 met15D0 ura3D0) strain background. A plasmid with yeast codon-optimized Dsup driven by doxycycline-inducible tet-on promoter system (NHB1054: untagged Dsup, or NHB1086: N-terminus tagged Dsup) was digested by NotI and integrated to the Leu2D0 locus of the background strain.

### Yeast viability experiments

Wild type and Dsup cultures were induced with 500 nM doxycycline for 8 hours to OD ∼0.6 and allowed to grow for up to 4 more hours in the presence of 2 µM hydrogen peroxide. Cells were collected by centrifugation before staining with Propidium Iodide (PI) dye (final concentration 6 µg mL^-1^) at room temperature for 2 minutes. Then 3 μL of each cell sample was transferred onto a glass slide and imaged with fluorescence microscopy.

### Fluorescence imaging

Microscopy experiments were performed using a Nikon Ti-E inverted fluorescence microscope with Perfect Focus, coupled with an EMCCD camera (Andor iXon X3 DU897). The light source is a spectra X LED system. Images were taken using a CFI plan Apochromat Lambda DM 60× oil immersion objective (NA 1.40 WD 0.13MM). The exposure and intensity setting for each channel were set as follows: Phase 50 ms, iRFP 10 ms at 10% lamp intensity with an EM Gain of 50, mRuby 50 ms at 10% lamp intensity with an EM Gain of 200. The EM Gain settings are within the linear range.

### FLIM-FRET imaging and analysis

HEK293T cells were imaged using a Leica SP8 FALCON confocal microscope equipped with a 63×/1.4 NA oil-immersion objective, a white-light laser (WLL), and a HyD SMD single molecule detector (Leica). For fluorescence imaging, EGFP was excited at 488 nm and collected with a 505–530 nm band-pass filter.

SYTO™ 59 fluorescence was exited at 610 nm and collected with a 630–670 nm band-pass filter. For FLIM imaging, EGFP was excited at 488 nm by using the WLL at a repetition rate of 40 MHz. Images were acquired, collecting maximal 1,000 photons per pixel. A region of interest (ROI) corresponding to the cell nucleus was selected per cell. Fast FLIM maps (512×512 pixels), fitting of fluorescence decay curves, and amplitude-weighted mean fluorescence lifetimes were calculated in LAS X (Lecia) software by fitting a bi-exponential decay model to the decay of each ROI. Graphs were plotted using Prism 8 (GraphPad).

### MD simulations

The sequence of Dsup residues 361-445 (Dsup for short) was from Uniprot entry P0DOW4: 361-PPRRSSRLTS SGTGAGSAPA AAKGGAKRAA SSSSTPSNAK KQATGGAGKA AATKATAAKS AASKAPQNGA GAKKKGGKAG GRKRK-445. Disordered conformations were generated using the TraDES method;^17^ three conformations with radii of gyration close to that expected^18^ for an IDP with 85 residues (*R*_g_ = 2.54 *N*^0.522^ = 25.8 Å for *N* = 85) were selected for three Dsup chains. Similarly to our previous study,^10^ a 35-base pair DNA was placed along the diagonal of a cubic box with a 120 Å side length; the three Dsup chains were randomly added to the box. Lastly water molecules along with eight Na^+^ ions were added to provide solvation and charge neutralization. The DNA, protein, and water force fields were Amber OL21,^19^ ff14SB,^20^ and OPC.^21^ The total number of atoms was 219,465.

The system was energy minimized for 5000 steps (half in steepest descent and half in conjugate gradient) using the sander module of AMBER22,^22^ followed by a 250 ps equilibration at constant NVT with a 1 fs time step, where the temperature was ramped from 0 to 300 K for the initial 100 ps and maintained at 300 K for the remaining 150 ps. Positional restraints were imposed on the Dsup chains and DNA with a force constant of 5 kcal mol^−1^ Å^−2^. Further equilibration was conducted without restraints for 1 ns with a 2 fs time step at constant NPT (300 K and 1 bar). Lastly, the simulations were continued in four replicates at constant NVT for 350 ns each with a 2 fs time step. All simulations were run on GPUs using pmemd.cuda.^23^ The cutoff for nonbonded interactions was 12 Å. Long-range electrostatic interactions were treated by the particle mesh Ewald method.^24^ The SHAKE algorithm^25^ was used to restrain bonds involving hydrogens. The temperature was maintained at 300 K by the Langevin thermostat with a damping constant of 1 ps^−1^, and the pressure was maintained at 1 bar by the Berendsen barostat.^26^

For DNA melting in the absence of Dsup, the 35-base pair DNA solvated by water and 68 Na^+^ ions (for charge neutralization) was energy minimized for 5000 steps and then equilibrated for 250 ps at constant NVT with a 1 fs time step, where the temperature was ramped from 300 to 500 K for the initial 100 ps and maintained at 500 K for the remaining 150 ps. The simulations were further continued in four replicates at constant NVT (500 K) for 200 ns. The simulations of DNA melting in the presence of Dsup were very similar, except that the initial snapshot was taken from the simulations at 300 K and energy minimization was skipped.

Snapshots were saved at 100 ps intervals, and analyses were done on the last 200 ns for the simulations at 300 K and for the entire trajectories for the simulations at 500 K. CPPTRAJ^27^ and in-house python and tcl scripts were used to calculate contacts within 3.5 Å between heavy atoms of Dsup and DNA. A base pair was considered melted when any hydrogen bond was broken (donor–acceptor distance > 3 Å).

### Statistics and reproducibility

All experiments were independently repeated as noted in the figure legends. Statistical analyses were performed using GraphPad Prism 8. For Gaussian data, Student’s t test was used for pairwise comparisons. For comparing three or more sets of data, ordinary one-way ANOVA followed by multiple comparisons test was performed. Statistical significance was defined *as P* < 0.05 with a 95% confidence interval. The number of cells analyzed (n cells) and number of independent experiments are reported in all figure legends. All time courses and scattered plots shown depict the mean ± standard error of the mean, unless otherwise noted.

## Data availability

The data that support the findings of this study is available within the main text and its Supplementary Information file. Source data is provided as Source Data file. Data is also available from the corresponding author upon request.

## Code availability

Custom ImageJ macros and MATLAB code used to analyze imaging data are available on GitHub.

## References

1. Bradford, A., & Weisberger, M. (2024a, February 23). What are Tardigrades and why are they nearly indestructible? LiveScience. https://www.livescience.com/57985-tardigrade-facts.html

2. Minguez-Toral, M., Cuevas-Zuviria, B., Garrido-Arandia, M. & Pacios, L.F. A computational structural study on the DNA-protecting role of the tardigrade-unique Dsup protein. Sci Rep 10, 13424 (2020).

3. Chavez, C., Cruz-Becerra, G., Fei, J., Kassavetis, G.A. & Kadonaga, J.T. The tardigrade damage suppressor protein binds to nucleosomes and protects DNA from hydroxyl radicals. Elife 8 (2019).

4. Aguilar, R., et al. Multivalent binding of the tardigrade Dsup protein to chromatin promotes yeast survival and longevity upon exposure to oxidative damage. Res Sq (2023).

5. Hashimoto, T. et al. Extremotolerant tardigrade genome and improved radiotolerance of human cultured cells by tardigrade-unique protein. Nat. Commun. 7, 12808 (2016).

6. Escarcega, R.D. et al. The Tardigrade damage suppressor protein Dsup promotes DNA damage in neurons. Mol Cell Neurosci 125, 103826 (2023).

7. Farrugia, G. & Balzan, R. Oxidative stress and programmed cell death in yeast. Front Oncol 2, 64 (2012).

8. Zandi, P. & Schnug, E. Reactive Oxygen Species, Antioxidant Responses and Implications from a Microbial Modulation Perspective. Biology (Basel) 11 (2022).

9. Reddy, V.P. Oxidative Stress in Health and Disease. Biomedicines 11 (2023).

10. Ahlawat, V., Dhiman, A., Mudiyanselage, H.E. & Zhou, H.X. Protamine-Mediated Tangles Produce Extreme Deoxyribonucleic Acid Compaction. J. Am. Chem. Soc. (2024).

11. Cremazy, F.G. et al. Imaging in situ protein-DNA interactions in the cell nucleus using FRET-FLIM. Exp. Cell Res. 309, 390–396 (2005).

12. Chidawanyika, T. & Supattapone, S. Hydrogen Peroxide-induced Cell Death in Mammalian Cells. J Cell Signal 2, 206–211 (2021).

13. Qi, L., Wu, X.C. & Zheng, D.Q. Hydrogen peroxide, a potent inducer of global genomic instability. Curr. Genet. 65, 913–917 (2019).

14. Das, A.T., Tenenbaum, L. & Berkhout, B. Tet-On Systems For Doxycycline-inducible Gene Expression. Curr. Gene Ther. 16, 156–167 (2016).

15. Kredel, S. et al. mRuby, a bright monomeric red fluorescent protein for labeling of subcellular structures. PLoS ONE 4, e4391 (2009).

16. Lin, W. et al. Light-gated integrator for highlighting kinase activity in living cells. Nat Commun 15, 7804 (2024).

17. Feldman, H.J. & Hogue, C.W.V. Probabilistic sampling of protein conformations: New hope for brute force? Proteins: Structure, Function, and Bioinformatics 46, 8–23 (2002).

18. Bernado, P. & Blackledge, M. A self-consistent description of the conformational behavior of chemically denatured proteins from NMR and small angle scattering. Biophys J 97, 2839–2845 (2009).

19. Zgarbová, M., Šponer, J. & Jurečka, P. Z-DNA as a Touchstone for Additive Empirical Force Fields and a Refinement of the Alpha/Gamma DNA Torsions for AMBER. Journal of Chemical Theory and Computation 17, 6292–6301 (2021).

20. Maier, J.A. et al. ff14SB: Improving the Accuracy of Protein Side Chain and Backbone Parameters from ff99SB. Journal of Chemical Theory and Computation 11, 3696–3713 (2015).

21. Izadi, S., Anandakrishnan, R. & Onufriev, A.V. Building Water Models: A Different Approach. The Journal of Physical Chemistry Letters 5, 3863–3871 (2014).

22. Case, D.A. et al. (University of California, San Francisco; 2023).

23. Salomon-Ferrer, R., Gotz, A.W., Poole, D., Le Grand, S. & Walker, R.C. Routine Microsecond Molecular Dynamics Simulations with AMBER on GPUs. 2. Explicit Solvent Particle Mesh Ewald. Journal of Chemical Theory and Computation 9, 3878–3888 (2013).

24. Essmann, U. et al. A Smooth Particle Mesh Ewald Method. J Chem Phys 103, 8577–8593 (1995).

25. Ryckaert, J.-P., Ciccotti, G. & Berendsen, H.J.C. Numerical integration of the cartesian equations of motion of a system with constraints: molecular dynamics of n-alkanes. Journal of Computational Physics 23, 327–341 (1977).

26. Berendsen, H.J.C., Postma, J.P.M., Vangunsteren, W.F., Dinola, A. & Haak, J.R. Molecular-Dynamics with Coupling to an External Bath. J Chem Phys 81, 3684–3690 (1984).

27. Roe, D.R. & Cheatham, T.E., 3rd PTRAJ and CPPTRAJ: Software for Processing and Analysis of Molecular Dynamics Trajectory Data. Journal of Chemical Theory and Computation 9, 3084–3095 (2013).

